# Localized ribosome access and distal tuning via the *Listeria prfA* RNA thermometer

**DOI:** 10.64898/2026.04.08.717274

**Authors:** Martin R. O’Steen, Jocelyn V. Chen, David H. Beier, Nils G. Walter, Sarah C. Keane

## Abstract

RNA thermometers (RNATs) are temperature-responsive structures in 5′ untranslated regions (UTRs) of bacterial messenger RNA (mRNA) that control translation by modulating ribosome access. The *Listeria monocytogenes prfA* RNAT represses translation of PrfA (positive regulatory factor A), the master virulence regulator, at ambient temperature and activates it near the human host temperature (∼37 °C) by modulating ribosome binding site (RBS) accessibility. However, the *prfA* RNAT shares no homology with known RNAT classes, and its unfolding mechanism remains unclear. Here, we used analytical ultracentrifugation and single-molecule kinetic analysis of RNA transient structure (SiM-KARTS) to map *prfA* RNAT unfolding. SiM-KARTS analysis demonstrates that thermal opening occurs predominantly at the RBS, while the upper helix of the RNAT hairpin remains largely folded at 37 °C. The RBS binding kinetics increases with temperature in parallel with translation output, establishing a quantitative link between structural unfolding and function. Mutations in the upper helix impair thermal regulation, indicating that this region tunes switching even as it stays structured at host temperature. Together, these data reveal a hierarchical unfolding pathway in which initial RBS opening triggers activation, whereas the upper helix remotely tunes temperature sensitivity.

**GRAPHICAL ABSTRACT:** 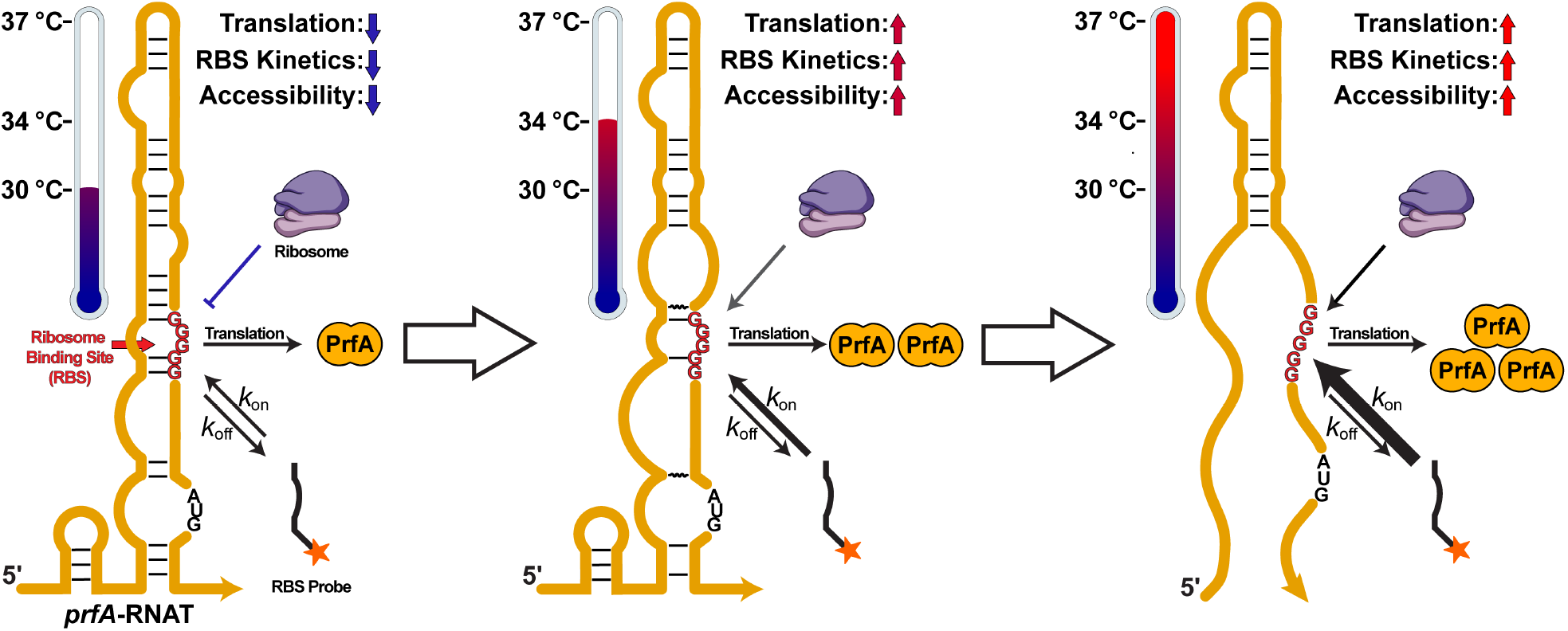

## INTRODUCTION

Non-coding RNA (ncRNA) elements in the 5′ untranslated region (UTR) of some bacterial messenger (m)RNAs are capable of modulating gene expression at both the transcriptional and translational levels [1–11]. 5′-UTR sequences that regulate translation in response to changes in temperature are known as RNA thermometers (RNATs) [1–3,12]. RNAT regulation relies on temperature-dependent structural changes that obscure or expose sequence elements required for translation, in particular the ribosome binding site (RBS) or start codon (AUG). This mechanism allows for rapid changes in mRNA translation in response to temperature changes, for example, upon cold or heat shock [4,5]. Additionally, RNATs are found in 5′-UTRs of some virulence genes and virulence gene transcription factors, enabling rapid adaptation to the host environment [1,6– 8].

*Listeria monocytogenes* is a serious food-borne pathogen that causes listeriosis, an illness with high rates of hospitalization and mortality for at risk populations [13]. The *prfA* RNAT in *L. monocytogenes* controls translation of positive regulatory factor A (PrfA), a master regulator of virulence gene expression [14]. The *prfA* RNAT is tuned to permit robust PrfA translation at host temperatures (∼37 °C) while nearly completely suppressing translation at ambient (or lower) temperatures [4,15]. Our previous work characterized the energetics of the *prfA* RNAT unfolding process, revealing a distinct two-transition unfolding profile [16]. Further mutational analysis demonstrated that internal loops and bridging helical domains influence the energetics and translatability of the *prfA* RNAT transcript. However, the structural dynamics underlying this translational control have remained unknown.

To correlate the translational regulation with structural changes within the *prfA* RNAT, we employed two solution characterization techniques; analytical ultracentrifugation (AUC) and single-molecule kinetic analysis of RNA transient structure (SiM-KARTS) [9–11,17,18]. AUC is a first-principles technique that can be used to study RNA folding and RNA interactions. While the structural resolution of AUC is lower than that of small angle X-ray scattering (SAXS), it is exceptionally sensitive to changes in RNA conformation as a function of solution conditions and has a tolerance for heterogeneity via the dynamic separation that occurs during the experiment. Complementarily, SiM-KARTS probes interrogate RNA structure fluctuations via the site-specific transient binding of small fluorescently labeled DNA probes. A total internal reflection fluorescence (TIRF) microscopy technique, SiM-KARTS facilitates the monitoring of structural fluctuations in a variety of solution conditions over long durations[18]. Quantification of the binding association constant (*k*_on_) provides a kinetic description of sequence accessibility and has been used to study RNA dynamics in riboswitches and other functional ncRNAs [9–11].

Our investigation of the *prfA* RNAT reveals that temperature-dependent unfolding begins at the RBS region, while the upper portion of the *prfA* RNAT remains structured. We further show that RNAT mutations that alter the stability of the RNA structure distal to the RBS significantly modulate accessibility of the RBS. These data are consistent with a model in which the influence of distal mutations is transmitted throughout the molecule, indicating that *prfA* RNAT function is influenced by the global, in addition to local, RNA structure.

## MATERIALS AND METHODS

### Analytical Ultracentrifugation

Sedimentation Velocity AUC experiments were conducted on Beckman Optima AUC at 26000 RPM at either ambient temperature (23 °C). 450-475 scans were collected with a scan interval time of 130-150s. All analysis was conducted in UltraScan III [19]. In brief, early traces are removed until a stable meniscus intensity is achieved with removal of extraneous baseline scans after *prfA* sedimentation. 2DSA analysis was performed with initial grid ranges of 1-7 s and 1-6 f/f_o_ with further refinement depending on the experimental condition. Following removal of time and radially invariant noise, noise and meniscus-corrected models underwent iterative modeling (10 iterations). Iterative model outputs were used to perform a GA analysis on the HPC located at the University of Texas Health Science Center [19,20]. Appropriate GA outputs were then utilized in subsequent Monte Carlo GA analysis to provide statistical measures of the derived parameters. More detailed procedures for collection and analysis are given in Table S1 and S2. Calculations were performed on the UltraScan LIMS cluster at the Bioinformatics Core Facility at the University of Texas Health Science Center at San Antonio.

### Fitting of the Magnesium compaction experiments

Frictional ratios were collected for a range of magnesium concentrations. The K_1/2,Mg_ was calculated in PRISM by fitting of data utilizing a log(inhibitor) vs response variable Hill slope equation which is shown below.

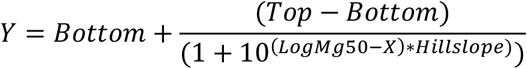

Top and bottom values were assigned based on the prfA RNAT’s frictional ratio at 0 mM Mg^2+^ (top) and the frictional ratio at 20 mM Mg^2+^ (bottom).

### Calculation of Radius of Hydration (R_h_)

Radius of hydration was calculated via the Stokes-Einstein Equation utilizing the diffusion coefficient and its associated statistical error from the MC-GA analysis.

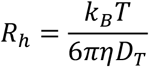

Where k_B_ is the Boltzmann constant, T is absolute temperature, η is the solvent viscosity and D_T_ is the translational diffusion coefficient.

### *In vitro* RNA transcription and purification

The *prfA* RNAT was generated through *in vitro* transcription as previously described [16]. Transcription reactions were carried out in 1X transcription buffer (40 mM Tris-Base pH 8.5, 15 mM dithiothreitol, 1 mM Spermidine, 0.01% Triton-X, 46.67 mM MgCl_2_), 2-5 mM rNTPs (weighted based on nucleotide composition of the RNA), 30-40 ng/µL DNA template, 0.2 units/mL yeast inorganic pyrophosphatase (New England Biolabs), 15 μM T7 RNA polymerase and 10% (v/v) dimethyl sulfoxide. Transcription reactions were incubated for 3 hours at 37 °C with shaking, then quenched with 1/6^th^ (v/v) 7 M urea and 250 mM ethylenediaminetetraacetic (EDTA) acid pH 8.5. The quenched transcription reaction was loaded onto a preparative-scale denaturing 12% polyacrylamide gel (National Diagnostics). Gels were visualized with UV shadowing and sections containing the RNA of interest were excised. RNA was extracted from the gel by crushing and soaking in TBE buffer (89 mM Tris, 89 mM boric acid, 2 mM EDTA, pH 8.3). The RNA was buffer exchanged first into 5 M sodium chloride to remove excess acrylamide and then into water using Amicon Centrifugal Filter Units (Millipore, Sigma). RNA integrity was checked on a 12% denaturing polyacrylamide gel. The purified RNA was incubated at 95 °C for 3 minutes then rapidly cooled at 4 °C to promote hairpin formation. RNA samples were aliquoted and stored at -70 °C to avoid freeze-thaw cycles. Sequences for all RNAs used are listed **S3 and S4**.

### *prfA* RNAT biotinylation

The 3′ end of purified *prfA* RNAT was modified with a single cytosine biotinylated nucleotide using the Pierce™ RNA 3′ End Biotinylation kit (Thermo Scientific, Waltham, MA USA), according to standard procedures.

### Fluorescent DNA oligonucleotides

Fluorescent DNA oligonucleotides were purchased from Integrated DNA Technologies. The sequences of the oligonucleotides used are listed in **Table S3**.

### Preparation for SiM-KARTS experiments

The SiM-KARTS construct was generated by mixing equimolar (1:1µM) solution of biotinylated *prfA* RNAT and capture probe in equal volumes. The solution was heated at 95 ºC for 3 minutes in water and slowly cooled to 4 °C over a period of 30 minutes. After capture probe annealing, the modified RNA was immobilized onto a PEG/PEG-biotin passivated coverslip using streptavidin. RNAs were then allowed to fold at experimental conditions for 15 minutes in fluorescence buffer (50 mM HEPES pH 7.5, 50 mM KCl, 1 mM MgCl_2_). Coverslips were washed with 3 volumes of fluorescence buffer to remove RNAs that lacked the biotin modification. SiM-KARTS experiments were carried out in fluorescence buffer supplemented with 5 mM protocatechuic acid, 50 nM protocatechuate dioxygenase and 2 mM Trolox to prevent fluorophore photobleaching. SiM-KARTS probes were added to concentrations of 50 nM for relevant experiments.

### SiM-KARTS data acquisition and analysis

All SiM-KARTS experiments were performed using a Nanoimager (ONI, Oxford, UK) using the 488 and 640 nm laser lines. The 488 nm laser line was used to localize immobilized *prfA* RNAT particles. The 640 nm laser line was used to visualize SiM-KARTS probe binding to the immobilized RNA. Two channel movies (488 and 640 nm) were collected for 1000 seconds.

Raw micrographs were analyzed using custom MATLAB (Mathworks) software as previously described [18]. Traces were manually selected and saved for kinetic analysis based on the following criteria: traces must contain at least 2 separate Cy5 fluorescent signals that were at least 2 times greater than background fluorescence, have single-step Cy5 fluorescent signals, and did not contain abnormalities. A two-state hidden Markov model (HMM) was applied to selected traces using the QuB software [21]. The “on” state dwell times (*τ*_bound_) approximated by the HMM corresponded to when the probe is bound to the RNAT, whereas “Off” state dwell times (*τ*_unbound_) corresponded to unbound SiM-KARTS probes. Dwell times for experimental replicates were compiled and fitted to single-exponential functions to determine the lifetimes of probe bound and unbound RNA states. *τ*_bound_ was converted to an association rate constant (*k*_on_) using equation

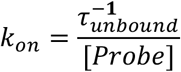

and *τ*_unbound_ was converted to a dissociation rate constant (*k*_off_) using equation

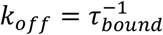

To convert kinetics to relative change in accessibility, the calculated kinetic constants were expressed as a ratio between the experimental condition and 30 °C.

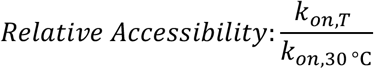

The two-way ANOVA with Tukey multiple comparisons test was used to determine statistical significance between SiM-KARTS probes at different temperatures.

### *In vitro* transcription/translation assays

350 ng of plasmid containing either prfA mutants or wild-type (WT), H4, or L5 were mixed with a final 1X PURExpress® solution from the PURExpress® In Vitro Protein Synthesis Kit (NEB #E6800), 0.7 µL RNaseOut™ Recombinant Ribonuclease Inhibitor (40 U/µL, Invitrogen #10777019), and 0.7 µL fluorescent lysine-charged tRNA from the FluoroTect™ GreenLys Translation Labeling System (Promega #L500A) for in-gel fluorescent visualization of protein expression. Reactions were incubated at 30, 34, and 37°C for 2 hours. 0.5 µL RNase A (Monarch® #T3018-2) was added to the residual translation reaction and incubated at 37 °C for 20 min. Following RNase A treatment, an SDS-PAGE buffer (50 mM Tris-HCl pH 6.8, 2% SDS, 10% glycerol, 1% β-mercaptoethanol, 12.5 mM EDTA) was added to 1X concentration, and the reactions were heated at 95 °C for 5 min. Electrophoresis was completed using house-cast gradient (8%-20%) SDS gels. The translation product containing BODFIPY FL Dye labeled lysine was visualized by fluorescent emission on an Amersham™ Typhoon™ biomolecular imager (Cytiva) using 488 nm laser excitation and a Cy2 525BP20 filter. Total protein visualization was completed via TCE staining. Conditions were normalized to the WT condition at each temperature. Variation in loading was accounted for by normalizing to a total stain protein band.

### Statistical analyses

Experimental data is presented as the mean of a minimum of three independent replicates and error bars represent the standard deviation of the replicates. Specific statistical tests are indicated in figure legends.

## RESULTS

### The *prfA* RNAT is well folded at physiological magnesium concentrations

We characterized the magnesium-induced compaction of the *prfA* RNAT by sedimentation velocity analytical ultracentrifugation (SV-AUC). RNA compaction was monitored by changes in the frictional ratio (f/f_0_), a parameter derived from the frictional coefficient. It relates to a molecule’s anisotropy, with f/f_0_ ranging from 1.0 to >4 for spherical and extended rod-like structures, respectively. Genetic algorithm Monte Carlo analysis was used to derive the f/f_0_ and all hydrodynamic values [20]. Fitting of the f/f_0_ at each magnesium concentration with the Hill equation (R^2^ = 0.995) yielded a K_1/2,Mg_ of 1.37 ± 0.16 mM Mg^2+^ (Fig. 1A). This magnesium concentration is physiologically relevant, as available cytosolic magnesium in prokaryotes is estimated to be ∼2.0 mM and reasonable given the sufficient monovalent cation concentration, 100 mM KCl, in our experimental conditions [22]. The reduction of the frictional ratios (2.35 to 2.0) paralleled a reduction in the radius of hydration (R_H_) from 44.5 to 39.7 Å (Fig. 1B). An overlay of the Monte Carlo modeling (Fig. 1C) demonstrates the expected correlation between sedimentation (s) and f/f_0_ with a narrowing of the structural ensemble at high magnesium concentrations. Full details on analysis and statistical outputs are available in Supplementary Tables S1 and S2 in the Supporting Material. These findings informed the best solution conditions for our kinetic analysis via SiM-KARTS.

**Figure 1.**
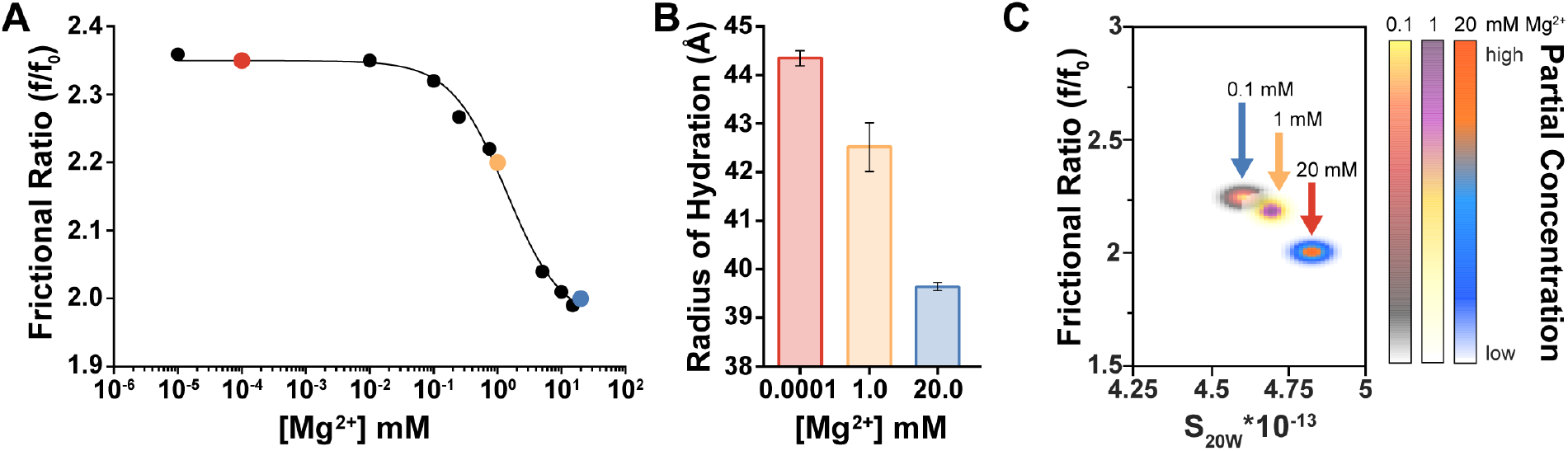
Compaction of the *prfA* RNAT upon addition of magnesium. (**A**) SV-AUC Mg^2+^ titrations reveal a reduction in frictional ratio, indicative of compaction of the RNA into a more spherical conformation. Red, yellow, and blue data points highlight 100 nM, 1 mM and 20 mM Mg^2+^, respectively. (**B**) Calculated radius of hydration (R_h_) at 100 nM, 1 mM and 20 mM Mg^2+^ demonstrate compaction across the titration. (**C**) A pseudo 3D overlay of the Monte Carlo Genetic Algorithm results at the indicated Mg^2+^ concentrations.

### Competitive Probes Modulate Binding Kinetics

SiM-KARTS facilitates the real-time monitoring of structural changes within an RNA molecule via transient interactions with short, fluorescently labeled oligonucleotide probes that bind to a surface-immobilized target RNA in a sequence-specific manner [18]. SiM-KARTS has been applied to a variety of riboswitches to characterize their dynamics in response to ligand binding [9–11]. We extended this technique to monitor local structural changes in the *prfA* RNAT in response to temperature. A schematic of the SiM-KARTS construct is shown in Fig. 2A. The target RNA is biotinylated at the 3′ end of the *prfA* RNAT allowing for immobilization on biotin passivated glass coverslip with streptavidin. An Alexa Fluor 488 (AF488) labeled locked nucleic acid (LNA) probe complementary to the 5′-end of the *prfA* RNAT was designed to localize immobilized RNAs. Single-site binding of the localization probe was confirmed via native gel electrophoresis. SiM-KARTS probes are designed to have melting temperatures (T_M_) near the experimental temperature, facilitating transient binding (Supplementary Table S3). A SiM-KARTS probe was designed to interrogate the structural dynamics at the ribosome binding site (RBS) region within the *prfA* RNAT (Fig. 2A, RBS probe). A full sequence diagram with annotated primer binding sites can be found in Fig. S2. Cumulative frequency plots for each condition depicting the k_on_ and k_off_ and number of molecules tracked are found in Supplementary Fig. S4.

**Figure 2.**
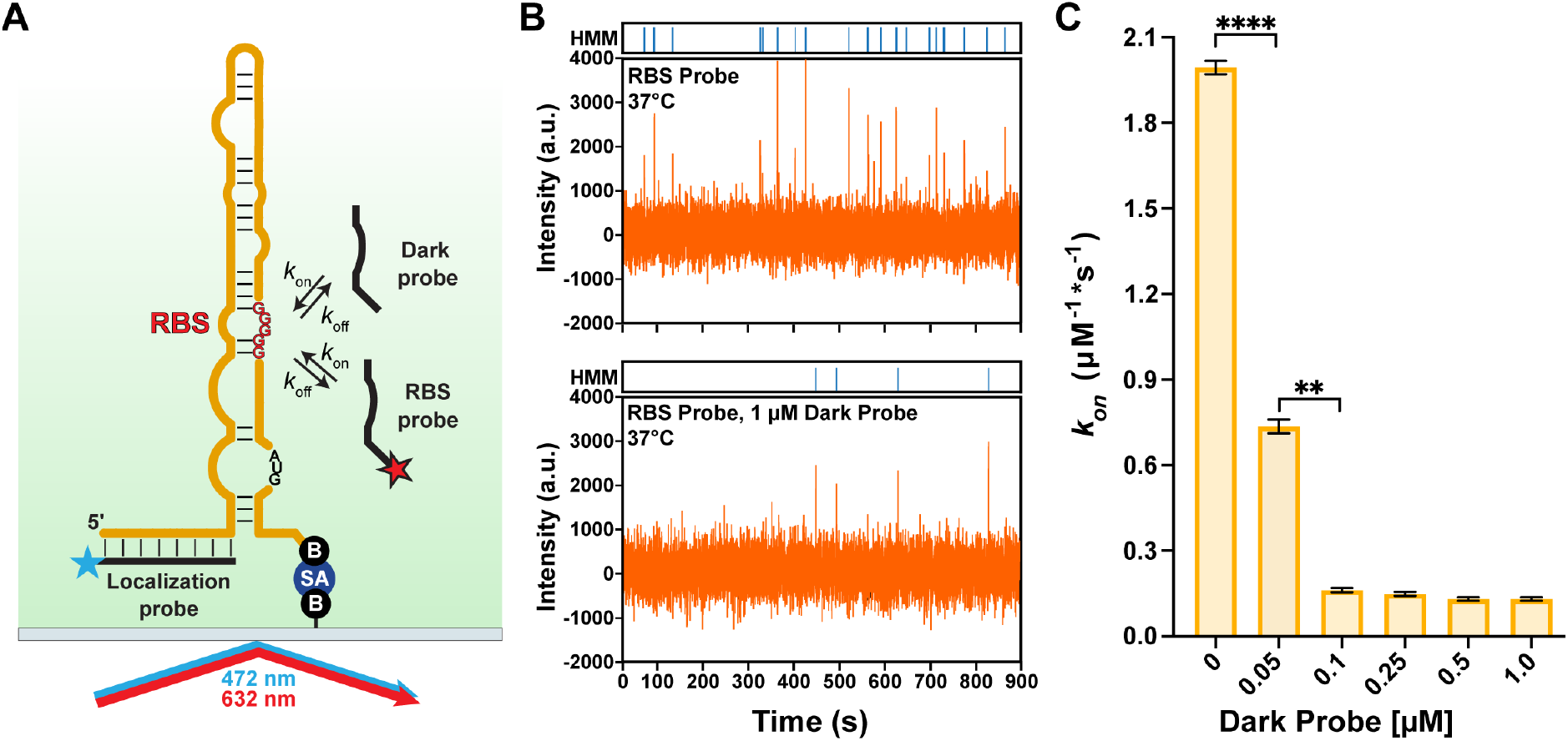
Competitive SiM-KARTS analysis of ribosome binding site (RBS) conformational dynamics. (**A**) Schematic of the SiM-KARTS construct showing the localization probe, laser excitation wavelengths and immobilization via biotin-streptavidin (B and SA) interaction on a biotin-PEG passivated cover slide. SiM-KARTS is total internal reflection technique, producing an evanescent field which excites fluorophore-labeled probes only when near the immobilized RNA. (**B**) Representative traces of the Cy5-labeled RBS probe in the absence (top) and presence of the dark probe (bottom). The binding frequency is greatly reduced as indicated by HMM idealization. (**C**) Kinetics analysis of the competitive SiM-KARTS experiment demonstrates reduced binding kinetics due to the presence of the unlabeled competitor probe for the RBS region. Statistical analysis was performed using Welch and Brown-Forsythe ANOVA tests. The data represent the kinetic model derived from a minimum 3 independent trials and shows the standard deviation.

The 5′-Cy5 labeled RBS probe is complementary to a short (8-10 nucleotide) region corresponding to the RBS, a region necessary for translation initiation. To validate our SiM-KARTS design, a competition experiment using a mixture of labeled and unlabeled RBS probe (dark probe, Fig. 2A) was developed. The RBS probe exhibited frequent binding to the RBS at 37 °C (Fig. 2B, top). Addition of the dark probe reduced the RBS probe binding frequency in a concentration dependent manner (Fig. 2B, bottom). Additional representative traces for competitive SiM-KARTS are found in Supplementary Fig. S3. At equimolar probe concentrations (50 nM), the dark probe reduced association binding kinetics more than 2-fold (Fig. 2C, Table 1).

**Table 1.** Summary of SiM-KARTS derived kinetics

| Competition Kinetic Values |  | Mutational Kinetic Values |  |  |
| --- | --- | --- | --- | --- |
| Dark Probe [$\mu\text{M}$] | $k_{\text{on}}$ ($\mu\text{M}^{-1}\text{s}^{-1}$) | Probe | Construct | $k_{\text{on}}$ ($\mu\text{M}^{-1}\text{s}^{-1}$) |
| 0.0 | $2.00 \pm 0.05$ | RBS Probe | *prfA-H4* | $2.41 \pm 0.03$ |
| 0.05 | $0.74 \pm 0.05$ | | *prfA-WT* | $2.00 \pm 0.05$ |
| 0.1 | $0.162 \pm 0.017$ | | *prfA-L5* | $0.89 \pm 0.03$ |
| 0.25 | $0.150 \pm 0.016$ | H4 Probe | *prfA-H4* | $1.66 \pm 0.17$ |
| 0.5 | $0.134 \pm 0.012$ | | *prfA-WT* | $0.308 \pm 0.01$ |
| 1.0 | $0.131 \pm 0.014$ | | *prfA-L5* | N/A |

At higher concentrations of dark probe (0.1 to 1 µM), labeled probe binding was significantly impaired, and no statistically significant changes in kinetics were observed. Taken together, these data demonstrate that our SiM-KARTS construct is valid, and that the RBS probe can be used to monitor RBS accessibility. We next employed a series of temperature dependent SiM-KARTS experiments to explore regional RNAT melting as a function of temperature.

### Temperature-dependent SiM-KARTS reveals the *prfA* RNAT unfolds first at the RBS

The translational control of RNATs relies on modulating RBS accessibility through temperature induced structural changes. To comprehensively probe the dynamic behavior of structural changes in the *prfA* RNAT, we employed a combination of temperature-dependent SiM-KARTS and *in vitro* translation assays [18,21]. We examined the structural dynamics of the *prfA* RNAT at two distinct regions, the RBS site and the H4 helix across physiologically relevant temperatures (Fig. 3A).

**Figure 3.**
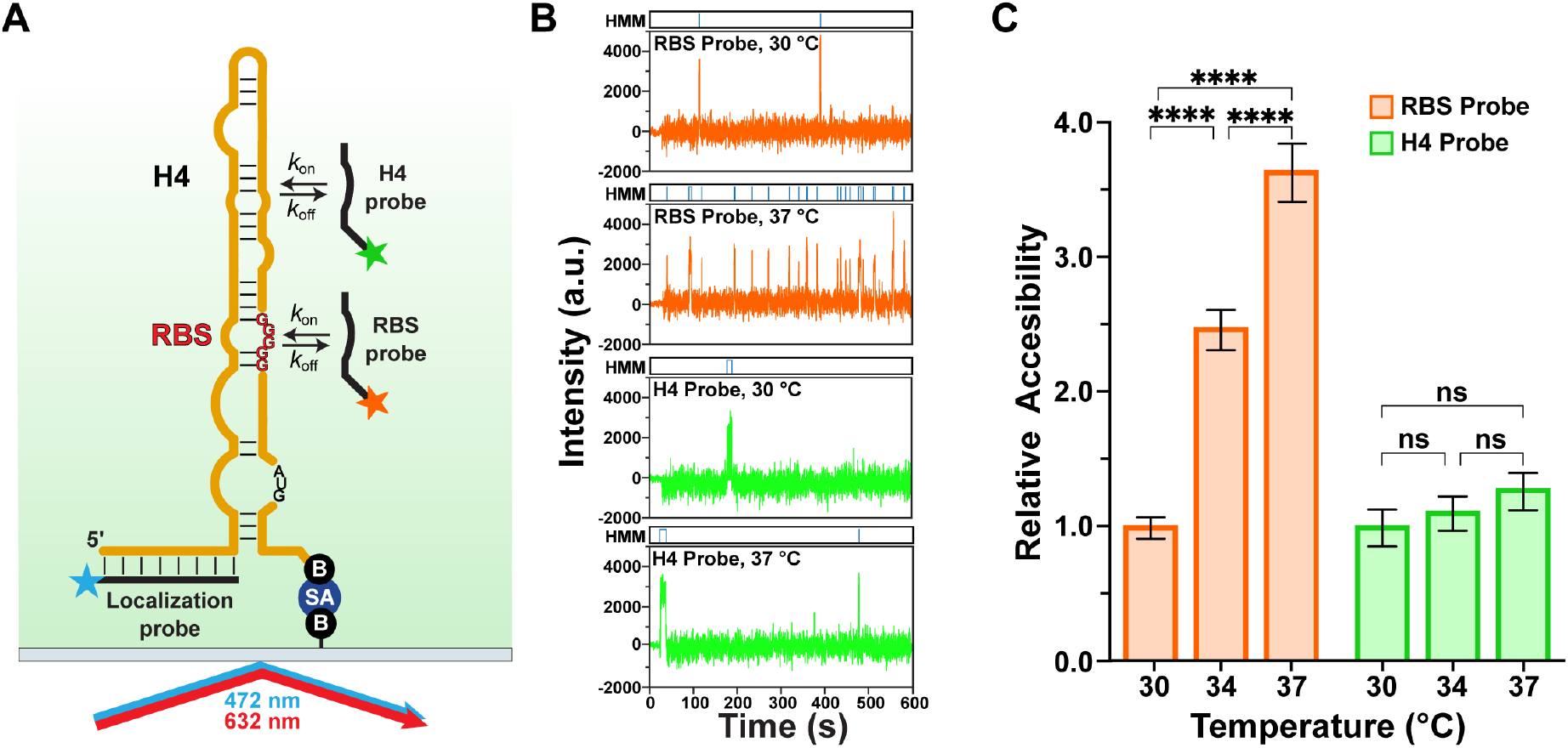
Probing the local dynamics of the *prfA* RNAT via SiM-KARTS. (**A**) Schematic of the SiM-KARTS construct. Independent experiments were conducted with a probe for either the RBS or H4 region. (**B**) Example SiM-KART traces for the RBS probe (top) and the H4 probe (bottom) at both 30 °C and 37 °C. HMM idealizations are shown above their respective traces. (**C**) The relative change in RBS (orange) and H4 (green) probe accessibility, normalized to the 30 °C condition for each probe. The H4 probe binding kinetics remain relatively constant, indicative of a well folded structure, however the RBS probe binding kinetics increase by a factor of 4 across the assayed temperatures. This change in accessibility indicates a loss of secondary structure in the region around the RBS. Statistical analysis was performed using Tukey multiple comparison with the two-way ANOVA. Data represent the kinetic model derived from a minimum 3 independent trials.

We quantified association rate constants for the RBS and H4 probes separately at 30, 34, and 37 °C (Fig. 3B). At 30 °C, the SiM-KARTS probes demonstrated similar association rate constants (*k*_on_) at both the RBS and H4 region of the *prfA* RNAT (0.168 ± 0.041 and 0.161 ± 0.035 µM^-1^*s^-1^, respectively; Fig. 3C). The slow association indicates that the *prfA* RNAT is well folded at 30 °C, restricting probe access at both the H4 helix and RBS site.

The kinetic constants for each probe at 34 and 37 °C were calculated and normalized to the 30 °C condition to provide relative changes in accessibility at the H4 helix and RBS site as a function of temperature (Fig. 3C). The H4 helix experienced only small changes in its limited accessibility across experimental temperatures (∼1.5-fold increase at 37 °C with respect to 30 °C), suggesting that the H4 helix remains folded even at translation permissible temperatures (37 °C). In contrast, the RBS probe rate constant increased dramatically over the same temperature range. The increased frequency of RBS probe binding at 34 (2.4-fold) and 37 °C (3.5-fold) demonstrates that the RBS region undergoes conformational changes, losing structure and becoming more accessible for probe binding at elevated temperatures. Cumulative frequency plots for both the H4 and RBS probe depicting k_on_ and k_off_ and number of molecules tracked are found in Supplementary Fig. S5.

### Translation output parallels RBS accessibility

We next determined whether changes in RBS accessibility correlate with translatability of the *prfA* mRNA. While *prfA* RNAT activity is well described at 20 °C and 37 °C (translation impermissible and permissible temperatures, respectively), other physiologically relevant temperatures and their influence on *prfA* translation remain underexplored[1,15,14]. Temperature dependent *in vitro* translation assays were conducted using plasmids containing either the prfA gene (containing the 5′-UTR and native coding sequence) or a dihydrofolate reductase (DHFR) gene (Fig. 4A). Translation reactions were supplemented with fluorescently labeled Lys-tRNA to enable in-gel analysis of the translation product (Fig. 4B). The fluorescence intensity of the protein product was quantified and normalized to the gene specific protein-levels at 37 °C for both PrfA and the DHFR control (Fig. 4B and 4C). The *dhfr* gene does not contain a 5′ RNAT element and, accordingly, the translation of DHFR was found not to be temperature dependent. Our studies revealed no statistical difference in DHFR protein levels across the experimental temperatures tested. PrfA translation at 30 °C is low, with a substantial increase at 37 °C, consistent with previous literature[23,15,16]. Interestingly, we found that the largest increase in translation occurred between 30 and 34 °C. These results are consistent with our observed increases in RBS accessibility, in which a 2.5-fold increase in RBS probe binding kinetics was observed between 30 °C and 34 °C (Fig. 4C).

**Figure 4.**
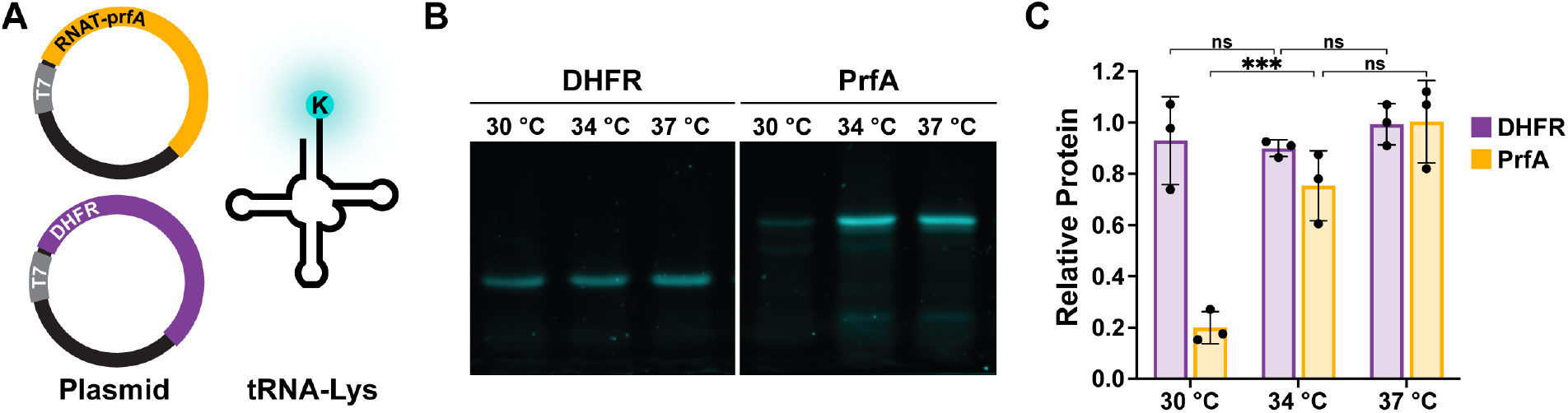
*In vitro* transcription/translation analysis reveals the temperature sensitivity of the *prfA* RNAT. (**A**) A schematic of the *in vitro* transcription/translation construct and labeled tRNA-Lys. A plasmid containing the *prfA* RNAT and PrfA coding sequence or the DHFR gene is added to the kit components and supplemented with fluorescently labeled tRNA-Lys to facilitate gel-based quantification of translation products. (**B**) Representative translation gel comparing the non-thermal regulated translation of DHFR against the *prfA* RNAT-regulated PrfA. (**C**) Quantification of *in vitro* translation results comparing DHFR to PrfA. The *prfA* construct demonstrates strong translation activation at 34 °C. The results represent the mean and standard deviation of 3 independent replicates.

### H4 helix mutations alter RBS accessibility and translatability

We next sought to determine whether mutations distal to the RBS influence the conformational dynamics of the *prfA* RNAT. Our previous work with the *prfA* RNAT developed a library of mutants that alter the underlying thermodynamics of the *prfA* RNAT and translatability of the downstream gene [16]. Two mutations, L5_stabilizing_ (L5) and H4_destabilizing_ (H4), located in the upper portion of the *prfA* RNAT hairpin, were selected for further temperature-dependent characterization (Fig. 5A). While the mutations are distal to the RBS, both affected the translational regulation of the *prfA* RNAT in *E. coli*, implying communication between the upper helical region and the central region of the *prfA* RNAT containing the RBS [16].

**Figure 5.**
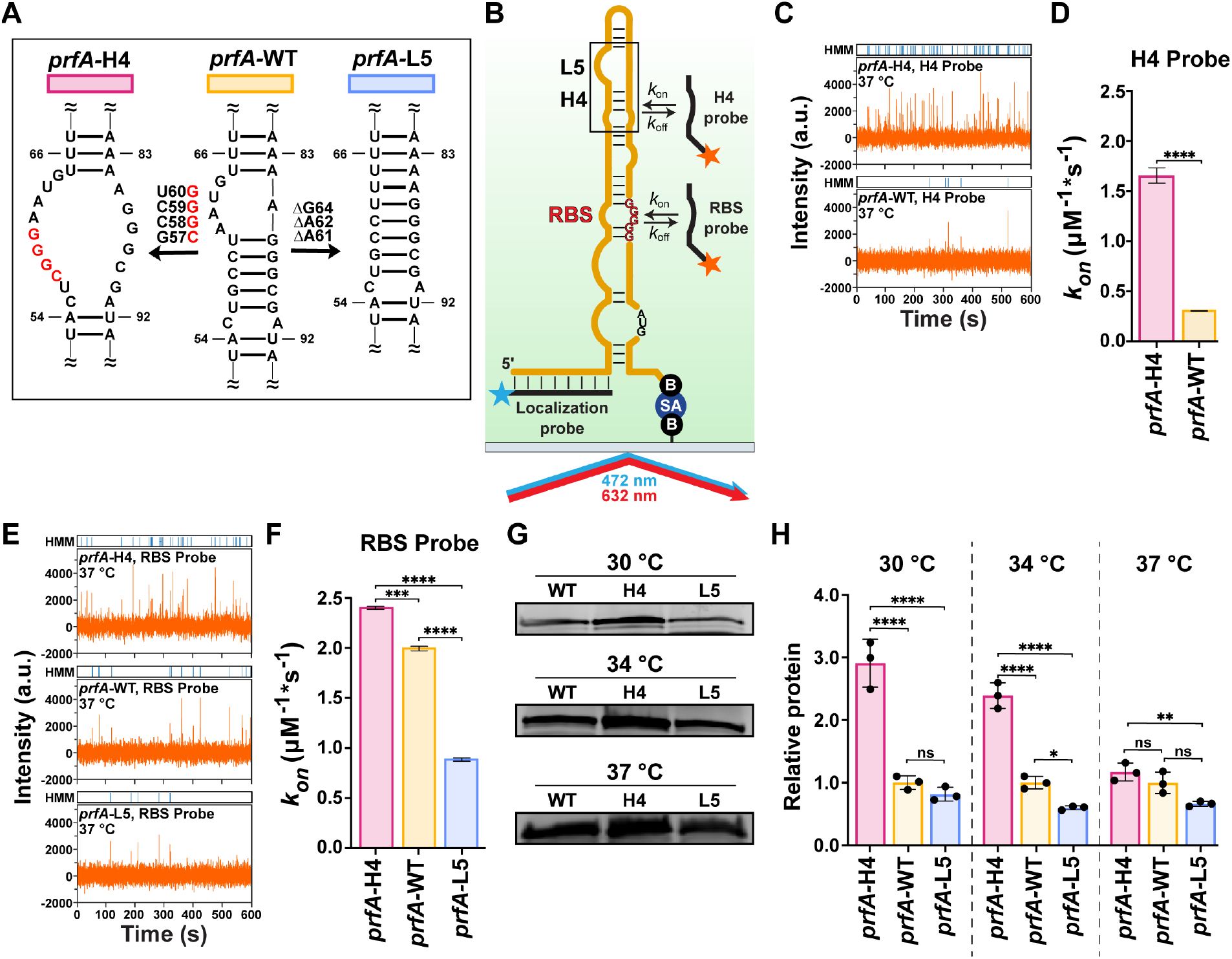
Mutations within the *prfA* RNAT alter RNA structure and impact temperature-regulated translation. (**A**) Secondary structure of the *prfA* RNAT (center) with the destabilizing (H4, left) and stabilizing (L5, right) mutations. (**B**) SiM-KARTS construct where the *prfA-WT*, -H4, and L5 RNATs were interrogated with RBS and H4 probes. (**C**) Example SiM-KART traces for *prfA*-H4 (top) and *prfA-WT* (bottom) with the H4 probe at 37 °C. HMM idealizations are shown above their respective traces. (**D**) Quantification of the H4 probe binding kinetics for *prfA*-H4 and *prfA-WT* revealed a higher frequency of binding to the *prfA*-H4 RNAT due to the destabilizing mutations. Data represents the derived kinetic models from a minimum of 3 replicates. Welch’s t test was used for statistical analysis. (**E**) Example SiM-KART traces for *prfA*-H4 (top) and *prfA-WT* (middle), and *prfA*-L5 (bottom) with the RBS probe at 37 °C. HMM idealizations are shown above their respective traces. (**F**) Quantification of the RBS probe binding kinetics for *prfA*-RNAT mutants revealed that distal mutations alter RBS accessibility. Data represent the kinetic models derived from a minimum of 3 independent experiments. A Welch and Brown-Forsythe one-way ANOVA was performed for statistical analysis. (**G**) PrfA expression levels for -WT, -H4, and -L5 constructs at 30 °C, 34 °C, and 37 °C. (**H**) Quantification of *in vitro* translation results comparing the -H4, -WT, and -L5 constructs at 30 °C, 34 °C, and 37 °C. All quantified product intensities were normalized to the -WT expression at a given temperature. Statistical analysis of relative protein expression used Tukey multiple within two-way ANOVA test and represent the mean of 3 independent trials.

We used SiM-KARTS to kinetically probe the *prfA*-H4 and *prfA*-L5 mutants at the H4 helical region and RBS, respectively (Fig. 5B-F). H4 kinetic analysis demonstrated mutation-induced structural differences in the H4 helical element for both *prfA*-H4 and *prfA*-L5 constructs. Destabilization of the H4 helical element, *prfA*-H4, resulted in much higher localized binding kinetics, reaching a nearly 6-fold increase in H4 probe binding kinetics relative to WT (*k*_on,*prfA*-H4_= 1.66 ± 0.077, *k*_on,*prfA-WT*_=0.3 ± 0.004 µM^-1^s^-1^) (Fig. 5C and 5D, Table 1). Stabilizing mutations in this region (*prfA*-L5) depressed H4 probe binding, precluding kinetic analysis at this position. Kinetic analysis at the RBS of the *prfA-WT* construct revealed an association kinetic constant of 2.00 ± 0.023 µM^-1^s^-1^ (Fig. 5E and 5F), comparable to those found for other functional RNA elements, such as riboswitches when analyzed by SiM-KARTS [9,10,17]. Both mutant RNATs (*prfA*-H4 and *prfA*-L5) exhibited altered RBS probe binding kinetics at 37 °C (Fig. 5E and 5F, Table 1), indicating that the stability of the upper portion of the RNAT influences global structure and RNAT function, consistent with our previous studies [16]. The *prfA*-H4 mutant’s lower thermodynamic stability was reflected by an increase in RBS probe binding kinetics compared to the WT RNAT (2.41 µM^-1^s^-1^ ± 0.014 and 2.00 µM^-1^s^-1^ ± 0.023 for the *prfA*-H4 and *prfA-WT* respectively) (Fig. 5E and 5F, Table 1). In contrast, the stabilizing mutation (*prfA*-L5) reduced RBS accessibility (0.894 µM^-1^s^-1^ ± 0.016 vs 2.00 µM^-1^s^-1^ ± 0.023, for *prfA*-L5 and *prfA-WT*, respectively) (Fig. 5E and 5F, Table 1), consistent with its reported increased stability and lower translatability. Cumulative frequency plots for each condition depicting the k_on_ and k_off_ and number of molecules tracked are found in Fig. S6.

Finally, we sought to establish how these distal mutations alter thermal regulation of *prfA* translation. The translation of the *prfA* constructs mirrors the RBS probe binding kinetics at 37 °C (Fig. 5G and 5H). Analysis of the *prfA*-L5 mutation demonstrated the expected lower translatability, 0.661 ± 0.04 relative expression to the *prfA-WT* construct (Fig. 5G and 5H), consistent with the difference in RBS kinetics at 37 °C. Even though the H4 mutation increases RBS accessibility, at 37 °C the expression of PrfA between *prfA-WT* and *prfA*-H4 is similar. The *prfA*-RNAT is naturally unfolded at 37 °C so that reducing the thermodynamic stability of the RNAT (*prfA*-H4) does not appreciably increase protein expression at this translation permissible temperature. In contrast, at lower temperatures (30 °C and 34 °C), the *prfA*-H4 construct had significantly higher translatability than *prfA-WT*. These findings support the notion that the H4 helix of the *prfA* RNAT participates in tuning the thermal response of the *prfA*-RNAT (Fig. 5G and 5H).

## DISCUSSION

RNA thermometers represent an efficient regulatory mechanism that allows bacteria to rapidly adapt to changes in the environmental temperature, which would be expected in the case of heat shock, cold shock, or host infection. Enteric pathogens face such challenges upon host consumption, and RNAT-based regulation has been exploited in the 5′ UTRs of virulence genes or virulence gene regulators, such as the *prfA* RNAT in *Listeria monocytogenes* [1,6,7]. The *prfA* RNAT regulates translation of positive regulatory factor A (PrfA), a key transcriptional activator for LM pathogenesis [14]. However, the *prfA* RNAT lacks sequence and structural homology with the canonical repression of heat shock gene expression (ROSE) and 4U RNAT families, raising the possibility that it represents a distinct mechanism of thermal regulation [24–27]. Our previous work characterized the unfolding of the *prfA* RNAT thermodynamically, revealing a non-cooperative unfolding profile. These findings were in contrast to unfolding studies of the 4U RNATs or the *cssA* RNAT, which exhibited more cooperative unfolding [16,27,28]. However, we were previously unable to assign these thermal transitions to specific regions within the RNAT structure. Mutational analysis of the *prfA* RNAT internal loop and helical elements distal to the RBS or start codon revealed a crucial role for these elements in tuning *prfA*’s unfolding and translatability. Here, we spatially refine the unfolding mechanism of the *prfA* RNAT by site-specific probing and mutagenesis. We demonstrate that RBS-distant mutations are transmitted to and modulate the accessibility of the *prfA* RNAT RBS.

AUC is well suited to interrogate RNA conformational changes, as hydrodynamic properties are sensitive to changes in molecular shape. We first performed a magnesium titration (0-20 mM Mg^2+^) and monitored changes in the frictional ratio of the *prfA* RNAT (Fig. 1A). Magnesium titration studies via AUC are typically applied to larger RNAs, such as long noncoding RNAs, whereas investigations of RNAs similar in size to the *prfA* RNAT (127 nt) are limited [29–33]. Our AUC analysis yielded a K_1/2,Mg_ of 1.37 ± 0.16 mM, closely aligned with prokaryotic cytosolic Mg^2+^ concentrations (1-2 mM), and similar to those of other natural RNA structures. Our solution conditions, 100 mM KCl, are sufficient to stabilize secondary structure of the *prfA* RNAT. The overall compaction, 5.5 Å, reflects the establishment of tertiary contacts or magnesium-dependent folds (Fig. 1B) [22].

We next sought to probe the regional conformational dynamics of the *prfA* RNAT using SiM-KARTS. While riboswitch structure and ligand binding have been studied using this single-molecule structure probing tool, it has not previously been used to study RNATs. Our analysis of the *prfA* RNAT demonstrates the applicability of this technique to a range of functional ncRNAs classes. RNATs inhibit or promote translation by sequestering or exposing translation sequence elements via temperature-induced structural changes. As an RNAT, *prfA* shares little sequence or structural homology to known RNAT families, leaving the mechanism of thermal regulation in question. Qualitatively, our kinetic analysis is similar to those of other functional ncRNA assayed by SiM-KARTS, even though a direct comparison across solution conditions and analytes must be made cautiously [9,10,18].

The *prfA* RNAT RBS and H4 helix exhibited unique kinetic profiles (Fig. 3B and 3C), consistent with distinct regional structural changes in response to temperature. The RBS probe exhibited a larger change in temperature-dependent SiM-KARTS probe binding kinetics, suggesting that the RBS is a structurally dynamic portion of the RNAT—consistent with secondary structure models predicting weak base pairing in the RBS vicinity. In contrast, the H4 helix is a stable helical element and exhibited only minimal changes across our experimental conditions. These data support a model wherein *prfA* RNAT unfolding is localized to the RBS region, while the H4 region of the RNAT remains well folded at elevated (host-like) temperature. Our kinetics analysis is reinforced by our temperature dependent *in vitro* translation assays where the largest gain in PrfA expression was observed between 30 °C and 34 °C (Fig. 4C). This increase in protein expression is well correlated with our measured kinetic probe-RBS binding increases, demonstrating a connection between RBS accessibility and molecular output.

We next sought to characterize the influence of distal mutations on the kinetics and translatability of PrfA. RNATs are sensitive to changes in sequence content, even mutations that maintain structural features (base pairing) of the RNAT can have profound effects on functional output and energetics of the structure [27]. Mutations in the 5′ portion of the *prfA* RNAT did not impact probe binding sites, allowing for direct kinetic comparisons between all constructs. We first used SiM-KARTS to probe localized structural changes in the H4 helix at 37 °C (Fig. 5C and 5D). As predicted, the destabilizing *prfA*-H4 mutation significantly increased the binding kinetics of the H4 probe compared to the *prfA-WT* RNAT, reflective of the destabilization and reduced secondary structure imparted by the mutations. In contrast, the stabilizing *prfA*-L5 mutation reduced the H4 probe binding frequency, consistent with increased stabilization and structure in the H4 helix.

SiM-KARTS probing of the RBS region of the *prfA*-H4 and *prfA*-L5 RNATs demonstrated that the distal mutations induce changes in global thermoregulation, a trait observed also in other RNATs such as the *htrA* RNAT [3]. Both *prfA*-H4 and *prfA*-L5 mutations altered RBS probe kinetics in the expected manner, with the destabilizing H4 mutation and the stabilizing L5 mutation increasing and decreasing RBS accessibility, respectively, relative to the *prfA-WT* RNAT (Fig 5E and 5F). These findings further demonstrate that the structure of the *prfA* RNAT RBS is not only controlled by local sequence context, but is the result of interplay between local and distal factors. As shown here, residual structure within H4 appears to fine tune RBS accessibility, and additionally may influence co-transcriptional folding. A similar hierarchical contribution is observed in ROSE RNATs, where upstream stem–loops (SL1–SL3) modulate the thermosensitive SL4 domain [26].

Comparative *in vitro* translation between the *prfA-WT*, -H4 and -L5 constructs demonstrated broad agreement with our kinetic results (Fig. 5G and 5H). While no statistically significant differences in translation were observed at 37 °C, the *prfA*-L5 construct had mildly reduced expression consistent with the lower RBS kinetics. At 37 °C the H4 and WT constructs had comparable RBS kinetics and protein expression levels, reflective of similar degrees of unfolding. At lower temperatures, 30 and 34 °C, the H4 construct demonstrated statistically elevated levels of PrfA expression. This significantly higher translatability at lower temperatures is indicative of a loss of thermoregulation due to mutation-induced structural changes. Our comparative analysis reinforces that the *prfA* RNAT structure is optimized for its thermal regulatory properties from both a sequence and structural perspective.

Collectively, these studies allow us to propose a model of the *prfA* RNAT unfolding mechanism. At elevated temperatures, structural changes first occur within the RBS region, liberating sequence elements necessary for translation initiation, and possibly further promoted upon 30S ribosomal subunit binding. The H4 helical domain of the *prfA* RNAT remains well folded at 37 °C, with its stability critical for thermal regulation, as demonstrated by the fact that the structural changes from both destabilizing and stabilizing H4 mutations are communicated throughout the entire molecule. These finding reinforce the notion that the sequence-structure relationship of RNATs is optimized down to the single nucleotide.

## DATA AND CODE AVAILABILITY

All study data and underlying code are available upon reasonable request from the authors.

## ACKNOWLEDGMENTS

This work was supported by the National Science Foundation (MCB-1942398 to S.C.K., DGE 1841052 to D.H.B.), and the National Institutes of Health (R35 GM138729-04S1 to S.C.K. and R35 GM131922 to N.G.W.). Calculations were performed on the UltraScan LIMS cluster at the Bioinformatics Core Facility at the University of Texas Health Science Center at San Antonio using multiple High Performance Computing clusters supported by NSF XSEDE Grant #MCB070038 (to Borries Demeler). We are grateful for Dr. Borries Demeler’s guidance for the AUC analysis of the prfA RNAT. We are grateful to members of Nils Walter lab, specifically Drs. Adrian Chauvier, Elizabeth Duran and Rosa Romero for training and guidance on SiM-KARTS assays. We thank members of the Keane Lab for helpful discussions.

## AUTHOR CONTRIBUTIONS

Conceptualization, S.C.K. and N.G.W.; Data collection, M.R.O, J.V.C, and D.H.B.; Data analyses, M.R.O.; Writing – original draft, M.R.O.; Writing – review and editing, M.R.O., D.H.B., J.V.C., N.G.W., and S.C.K.

## DECLARATION OF INTEREST

The authors declare no competing interests.

## Notes

### Competing Interest Statement

The authors have declared no competing interest.

## REFERENCES

1. J Johansson P Mandin A Renzoni et al. An RNA Thermosensor Controls Expression of Virulence Genes in Listeria monocytogenes. Cell 2002;110:551–561. 10.1016/S0092-8674(02)00905-4

2. RP Barnwal E Loh KS Godin et al. Structure and mechanism of a molecular rheostat, an RNA thermometer that modulates immune evasion by Neisseria meningitidis. Nucleic Acids Res. 2016;gkw584. 10.1093/nar/gkw584

3. EK Choi KA Ulanowicz YAH Nguyen et al. SHAPE analysis of the *htrA* RNA thermometer from *Salmonella enterica*. RNA 2017;23:1569–1581. 10.1261/rna.062299.117

4. AM Giuliodori F Di Pietro S Marzi et al. The cspA mRNA Is a Thermosensor that Modulates Translation of the Cold-Shock Protein CspA. Mol. Cell 2010;37:21–33. 10.1016/j.molcel.2009.11.033

5. F Narberhaus. Translational control of bacterial heat shock and virulence genes by temperature-sensing mRNAs. RNA Biol. 2010;7:84–89. 10.4161/rna.7.1.10501

6. SM Brewer C Twittenhoff J Kortmann et al. A Salmonella Typhi RNA thermosensor regulates virulence factors and innate immune evasion in response to host temperature. PLOS Pathog. 2021;17:e1009345. 10.1371/journal.ppat.1009345

7. ER Murphy J Roßmanith J Sieg et al. Regulation of OmpA Translation and Shigella dysenteriae Virulence by an RNA Thermometer. Infect. Immun. 2020;88:e00871–19. 10.1128/IAI.00871-19

8. GG Weber J Kortmann F Narberhaus et al. RNA thermometer controls temperature-dependent virulence factor expression in *Vibrio cholerae*. Proc. Natl. Acad. Sci. 2014;111:14241–14246. 10.1073/pnas.1411570111

9. A Chauvier P Ajmera R Yadav et al. Dynamic competition between a ligand and transcription factor NusA governs riboswitch-mediated transcription regulation. Proc. Natl. Acad. Sci. 2021;118:e2109026118. 10.1073/pnas.2109026118

10. A Chauvier SS Dandpat R Romero et al. A nascent riboswitch helix orchestrates robust transcriptional regulation through signal integration. Nat. Commun. 2024;15:3955. 10.1038/s41467-024-48409-8

11. RA Romero VA Reed A Chauvier et al. Single-molecule investigation of the glycine tandem riboswitch and its role in gene regulation. Biophys. J. 2024;123:454a. 10.1016/j.bpj.2023.11.2763

12. I Borovská C Zhang S-LJ Dülk et al. Identification of conserved RNA regulatory switches in living cells using RNA secondary structure ensemble mapping and covariation analysis. Nat. Biotechnol. 2025;1–13. 10.1038/s41587-025-02739-0

13. Y Su A Liu and M-J Zhu. Mapping the landscape of listeriosis outbreaks (1998–2023): Trends, challenges, and regulatory responses in the United States. Trends Food Sci. Technol. 2024;154:104750. 10.1016/j.tifs.2024.104750

14. M Scortti HJ Monzó L Lacharme-Lora et al. The PrfA virulence regulon. Microbes Infect. 2007;9:1196–1207. 10.1016/j.micinf.2007.05.007

15. E Loh F Memarpour K Vaitkevicius et al. An unstructured 5′-coding region of the prfA mRNA is required for efficient translation. Nucleic Acids Res. 2012;40:1818–1827. 10.1093/nar/gkr850

16. H Zhang I Hall AJ Nissley et al. A Tale of Two Transitions: The Unfolding Mechanism of the prfA RNA Thermosensor. Biochemistry 2020;59:4533–4545. 10.1021/acs.biochem.0c00588

17. S Ray SS Dandpat S Chatterjee et al. Precise tuning of bacterial translation initiation by non-equilibrium 5′-UTR unfolding observed in single mRNAs. Nucleic Acids Res. 2022;50:8818–8833. 10.1093/nar/gkac635

18. A Chauvier J Cabello-Villegas and NG Walter. Probing RNA structure and interaction dynamics at the single molecule level. Methods 2019;162–163:3–11. 10.1016/j.ymeth.2019.04.002

19. B Demeler. UltraScan: a comprehensive data analysis software package for analytical ultracentrifugation experiments. Mod. Anal. Ultracentrifugation Tech. Methods 2005;10:210–229.

20. E Brookes and B Demeler. Genetic algorithm optimization for obtaining accurate molecular weight distributions from sedimentation velocity experiments. Anal. Ultracentrifugation VIII 2006;33–40.

21. F Qin and L Li. Model-Based Fitting of Single-Channel Dwell-Time Distributions. Biophys. J. 2004;87:1657–1671. 10.1529/biophysj.103.037531

22. C Hurwitz and CL Rosano. The intracellular concentration of bound and unbound magnesium ions in Escherichia coli. J. Biol. Chem. 1967;242:3719–3722.

23. E Loh O Dussurget J Gripenland et al. A trans-Acting Riboswitch Controls Expression of the Virulence Regulator PrfA in Listeria monocytogenes. Cell 2009;139:770–779. 10.1016/j.cell.2009.08.046

24. S Chowdhury C Ragaz E Kreuger et al. Temperature-controlled Structural Alterations of an RNA Thermometer. J. Biol. Chem. 2003;278:47915–47921. 10.1074/jbc.M306874200

25. S Halder and M Bansal. The effect of mutation in the stem of the MicroROSE thermometer on its thermosensing ability: insights from molecular dynamics simulation studies. RSC Adv. 2022;12:11853–11865. 10.1039/D2RA00169A

26. EA Jolley KM Bormes and PC Bevilacqua. Upstream Flanking Sequence Assists Folding of an RNA Thermometer. J. Mol. Biol. 2022;434:167786. 10.1016/j.jmb.2022.167786

27. J Rinnenthal B Klinkert F Narberhaus et al. Modulation of the stability of the Salmonella fourU-type RNA thermometer. Nucleic Acids Res. 2011;39:8258–8270. 10.1093/nar/gkr314

28. J Rinnenthal B Klinkert F Narberhaus et al. Direct observation of the temperature-induced melting process of the Salmonella fourU RNA thermometer at base-pair resolution. Nucleic Acids Res. 2010;38:3834–3847. 10.1093/nar/gkq124

29. D Costantino and JS Kieft. A preformed compact ribosome-binding domain in the cricket paralysis-like virus IRES RNAs. RNA 2005;11:332–343. 10.1261/rna.7184705

30. CE Dann CA Wakeman CL Sieling et al. Structure and Mechanism of a Metal-Sensing Regulatory RNA. Cell 2007;130:878–892. 10.1016/j.cell.2007.06.051

31. P Saha S Patel RCa Tavares et al. Structural basis for the function of long noncoding RNA Pnky in neural stem cells. 2025; 10.1101/2025.09.01.671568

32. S Somarowthu M Legiewicz I Chillón et al. HOTAIR Forms an Intricate and Modular Secondary Structure. Mol. Cell 2015;58:353–361. 10.1016/j.molcel.2015.03.006

33. F Liu S Somarowthu and AM Pyle. Visualizing the secondary and tertiary architectural domains of lncRNA RepA. Nat. Chem. Biol. 2017;13:282–289. 10.1038/nchembio.2272

