## Supplementary Data for "Localized ribosome access and distal tuning via the *Listeria prfA* RNA thermometer"

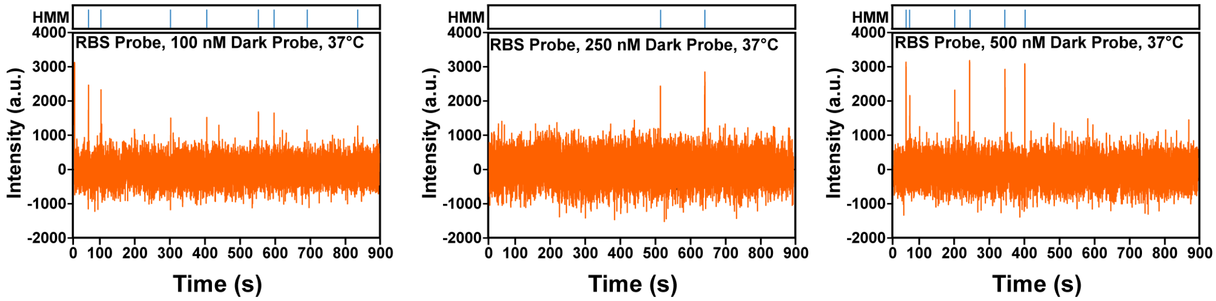


**Figure S1. Representative traces for 100, 250 and 500 nM SiM-KARTS competition assays.** The hidden Markov model (HMM) idealization for each condition is shown above. All data was collected at 37°C using the same conditions described in the main text.

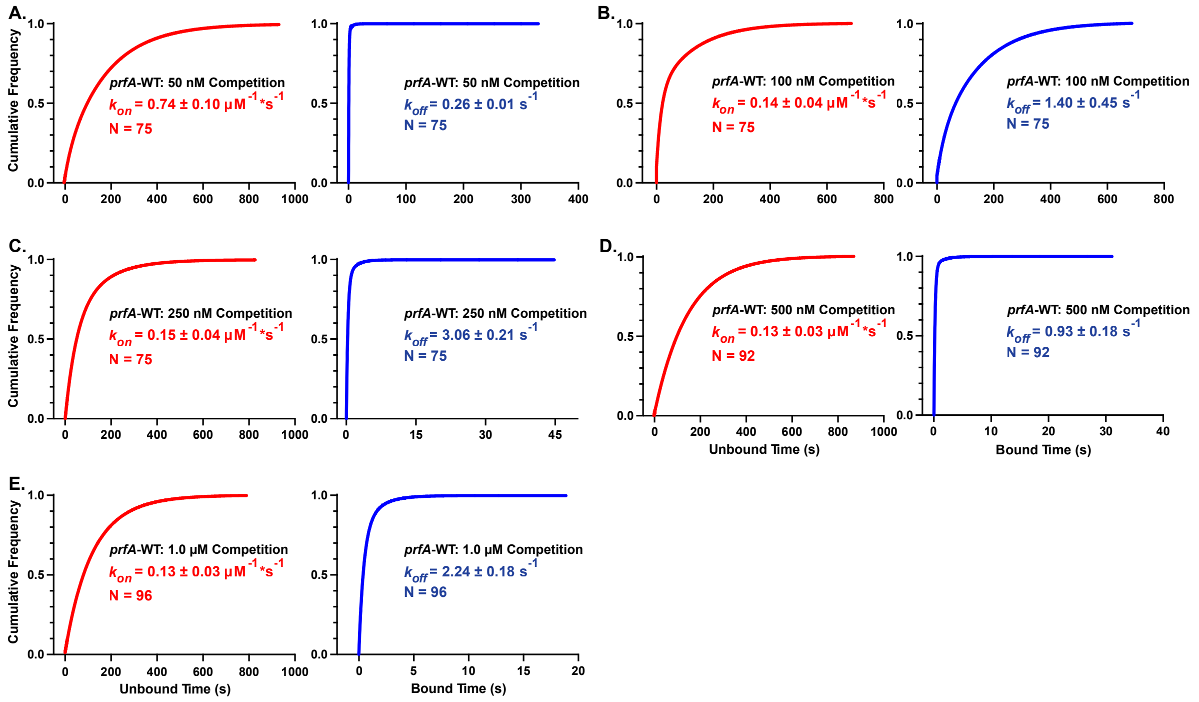


**Figure S2. Cumulative probability plots of unbound and bound dwell times for competition SiM-KARTs.** Association rate constants ($k_{on}$, red) and dissociation constants ($k_{off}$, blue) are given for the corresponding experimental condition. Number of molecules tracked for each experiment are shown by N. Values correspond to those depicted in Fig. 2 with (A) 50 nM, (B) 100, (C) 250, (D) 500 nM and (E) 1 μM. All fits achieved an $R^2$ (adj) of 0.999 except for 100 nM condition (B), whose $R^2$ (adj) was 0.997. All data was collected at 37°C using the same conditions described in the main text.

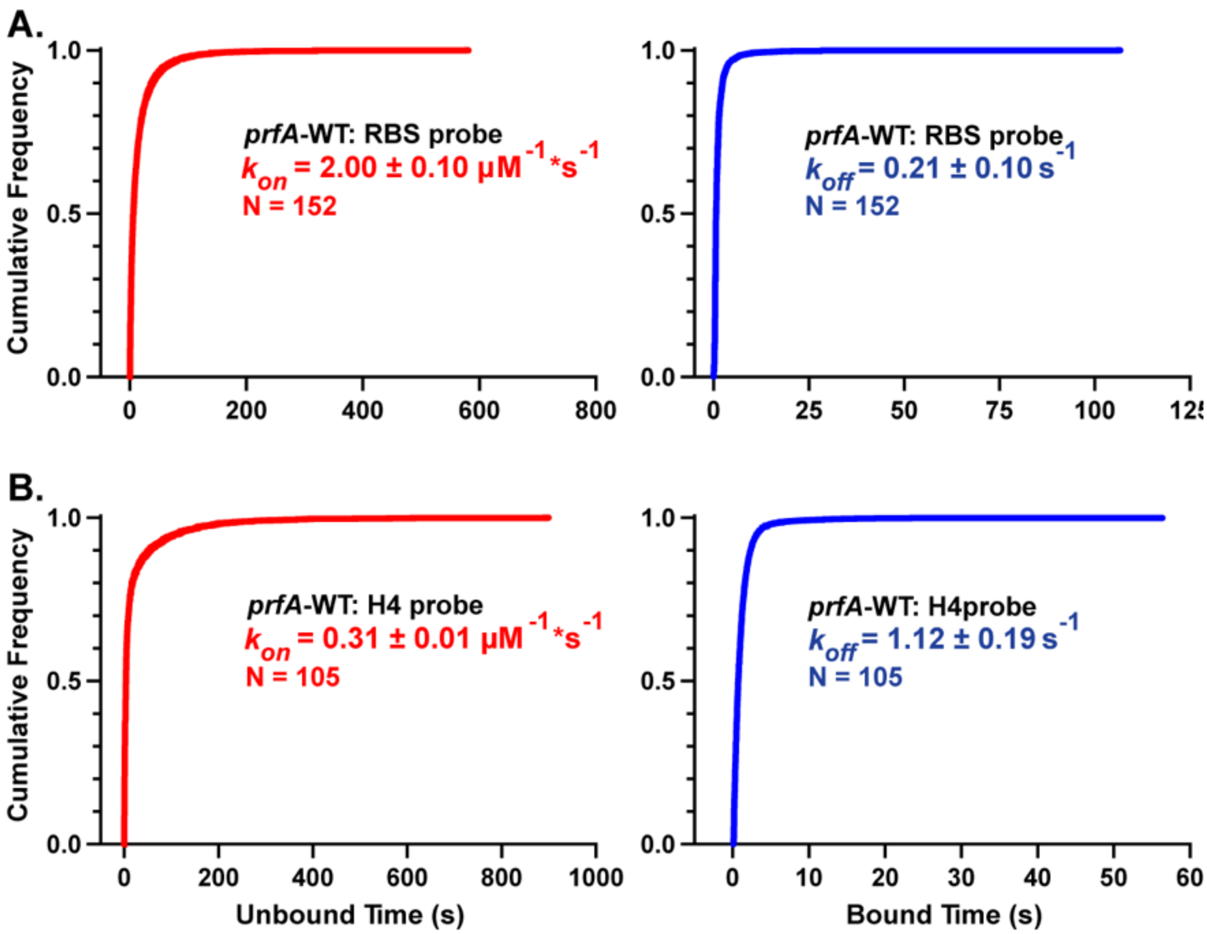


**Figure S3. Cumulative probability plots of unbound and bound dwell times for *prfA*-WT SiM-KARTs RBS and H4 analysis.** Association rate constants ($k_{on}$, red) and dissociation constants ($k_{off}$, blue) are given for the corresponding experimental condition. Number of molecules tracked for each experiment are shown by N. All data was collected at 37°C using the same conditions described in the main text. (A) The cumulative probability plots for the RBS Probe with *prfA*-WT at 37 °C. (B) The cumulative probability plots for the H4 probe with *prfA*-WT at 37 °C. $R^2$ adjusted values were 0.993 and 0.997 for A and B, respectively.

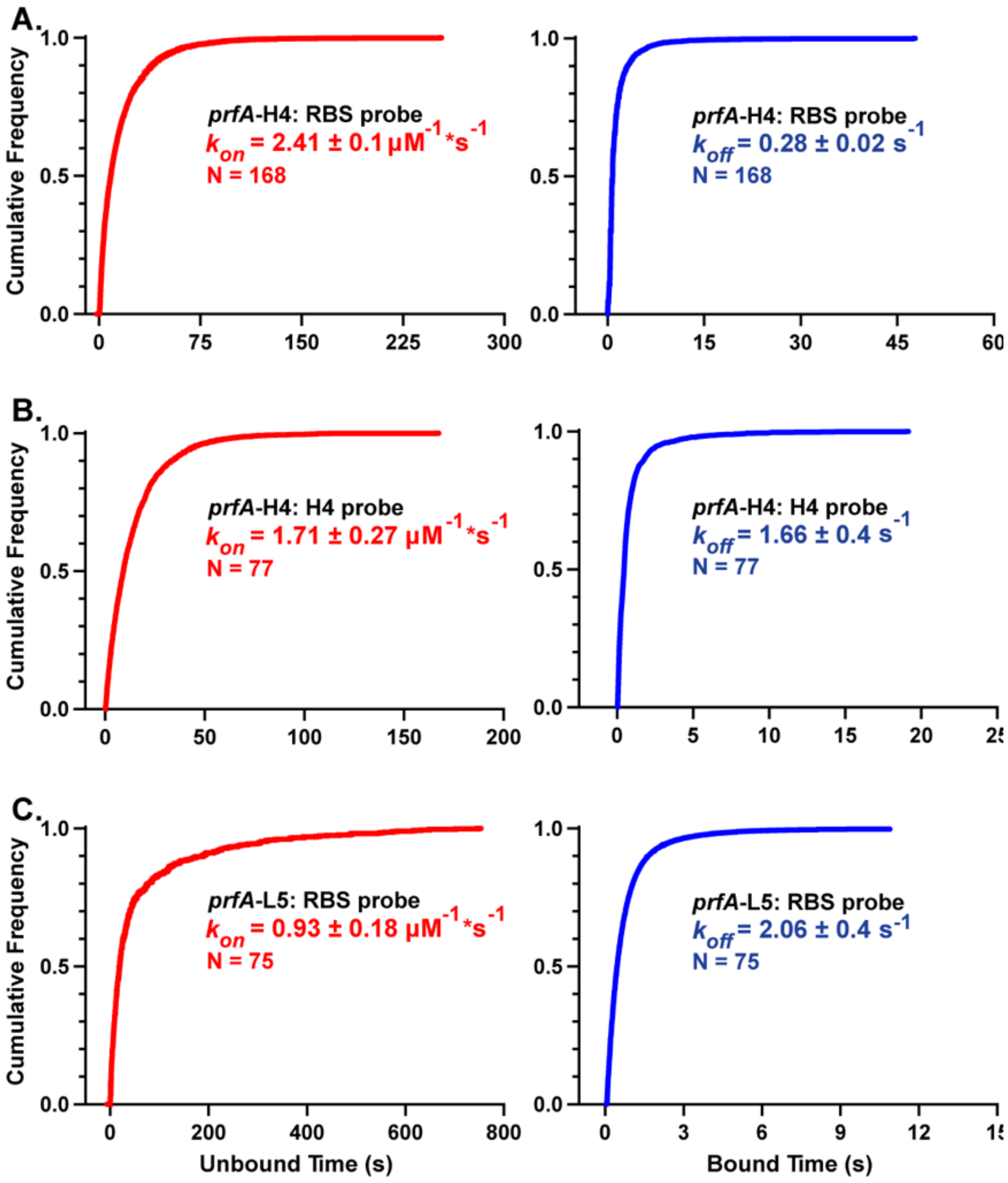


**Figure S4: Cumulative probability plots of unbound and bound dwell times for mutant SiM-KARTs analysis.** Association rate constants ($k_{on}$, red) and dissociation constants ($k_{off}$, blue) are given for the corresponding experimental condition. Number of molecules tracked for each experiment are shown by N. All data was collected at 37°C using the same conditions described in the main text. The cumulative probability plots for *prfA*-H4 RNAT with the RBS probe (A) and H4 probe (B). The cumulative probability plots for *prfA*-L5 RNAT with the RBS probe are shown in (C). All fits achieved an $R^2$ (adj) of 0.997 except for *prfA*-L5 (C), whose $R^2$ (adj) was 0.993. All data was collected at 37°C using the same conditions described in the main text.

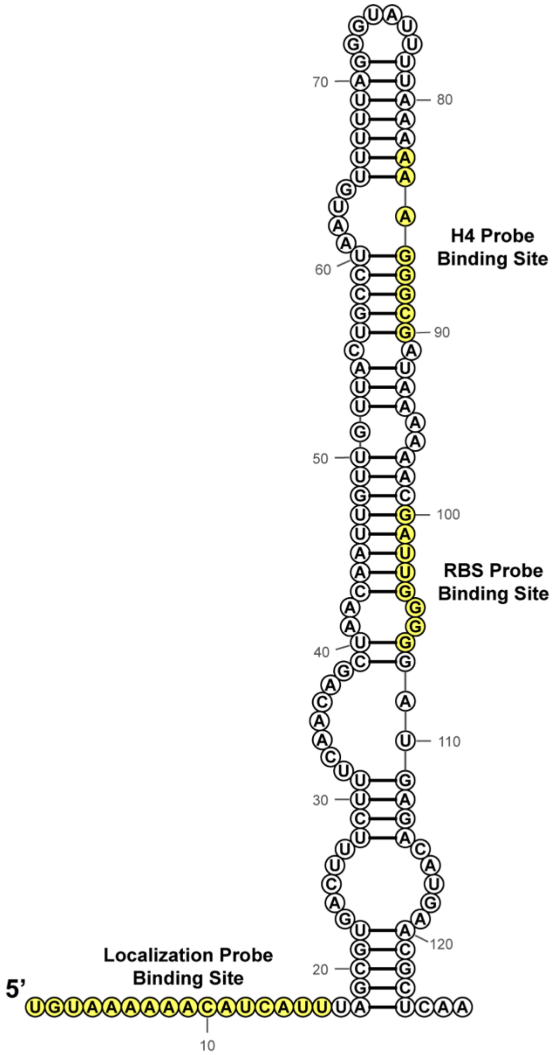


**Figure S5. Secondary structure sequence diagram of the *prfA*-WT RNAT.** The positions and complementary sequences to the H4, RBS and Localization probes are indicated by yellow fill and denoted. Primer binding sites are conserved in the *prfA*-H4 and *prfA*-L5 constructs as mutations are found between positions 57-64 on the 5' side of the predicted hairpin.

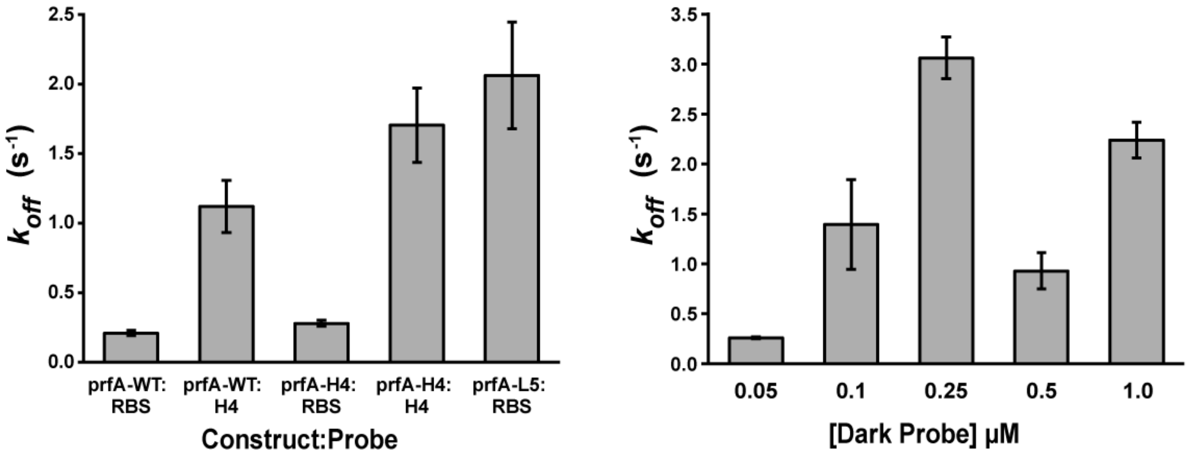


**Fig S6. Dissociation rate constants of mutant and competitive SiM-KARTs analysis.** (Right) Dissociation rate constants ($k_{off}$) of *prfA*-WT, *prfA*-H4, and *prfA*-L5 with the RBS and H4 Probe. (Left) Dissociation rate constants ($k_{off}$) of *prfA*-WT in the competition SiM-KARTs assay. The corresponding probability plots are given in Fig S2,3, and 4.

**Table S1. Hydrodynamic parameters of the *prfA*-WT RNAT.**

| $MgCl_2$ [mM] | Frictional Ratio ($f/f_0$) | $MW_{app}$ (kDa)[a] | Diffusion | Viscosity (cP) |
|---|---|---|---|---|
| 0.00 | 2.36 ± 0.07 | 39.9 | 4.87 ± 0.08 | 1.00521 |
| $10^{-4}$ | 2.35 ± 0.02 | 39.7 | | |
| $10^{-2}$ | 2.35 ± 0.01 | 40.3 | | |
| 0.10 | 2.32 ± 0.01 | 40.6 | | |
| 0.25 | 2.27 ± 0.01 | 40.3 | | |
| 0.75 | 2.22 ± 0.01 | 40.1 | 5.12 ± 0.06 | 1.00532 |
| 1.00 | 2.19 ± 0.02 | 40.6 | | |
| 5.00 | 2.04 ± 0.01 | 40.7 | | |
| 10.00 | 2.01 ± 0.02 | 40.5 | | |
| 15.00 | 2.00 ± 0.05 | 39.3 | | |
| 20.00 | 2.01 ± 0.003 | 41.5 | 5.39 ± 0.01 | 1.01419 |

a. Model derived molecular weights for AUC analysis of the *prfA*-RNAT (40.201 kDa)
b. Diffusion Coefficients used to calculate radius of hydration ($R_H$) in Figure 1B. Units are $\mathbf{10^{-7} * cm^2 * s^{-1}}$
c. Viscosity of solutions used to calculate radius of hydration ($R_H$) in Figure 1B. Values are from UltraScan-III solution module and given in centipoise (cP).

**Table S2. AUC analysis editing and fit of the *prfA*-WT RNAT.**

| Meniscus (cm) | Data Start (cm) | Data End (cm) | Total Scans | GA-MC[a.] | RMSD (* $10^{-3}$) |
|---|---|---|---|---|---|
| 6.073 | 6.130 | 7.017 | 450 | 96 | 2.41 |
| 6.043 | 6.111 | 7.022 | 500 | 96 | 2.07 |
| 5.956 | 6.021 | 7.01 | 473 | 96 | 2.12 |
| 6.058 | 6.145 | 7.012 | 500 | 48 | 2.08 |
| 6.063 | 6.144 | 7.01 | 450 | 48 | 2.51 |
| 6.081 | 6.130 | 7.03 | 450 | 96 | 2.49 |
| 6.066 | 6.127 | 7.011 | 500 | 48 | 2.4 |
| 6.075 | 6.110 | 7.014 | 450 | 96 | 2.49 |
| 6.067 | 6.138 | 7.017 | 400 | 96 | 2.42 |
| 6.075 | 6.110 | 7.014 | 400 | 72 | 2.42 |
| 6.036 | 6.102 | 7.05 | 450 | 96 | 2.44 |

a. Genetic Algorithm Monte Carlo Iterations

**Table S3. Probe sequences and experimental melting temperatures for SiM-KARTs analysis.**

| Probe | Sequence (5' to 3')[a,] | $Tm_{exp}$ (ºC)[b.] |
|---|---|---|
| H4 | /5Cy5/TACGCCCTTT | 31.9 |
| RBS | /5Cy5/GACCCAATC | 30.3 |
| Localization | ATT+G+A+T+G+T+TT+T+TA+C+A/3AlexF488N/ | 69.7 |

a. + indicates inclusion of LNA nucleotide

b. Melting temperatures were calculated at experimental conditions (50 mM $M^+$ and 1 mM $MgCl_2$) using DNA probe and RNA target.

### Table S4: *prfA* RNAT sequences used in this study.

| Construct | Sequence (5' to 3')[a,] |
|---|---|
| *prfA*-WT | UGUAAAAAACAUCAUUUAGCGUGACUUUCUUUCAACAGCUAACAAUUGUUGUUACUGCC UAAUGUUUUUAGGGUAUUUUAAAAAAGGGCGAUAAAAAACGAUUGGGGGAUGAGACAUG AACGCUCAA |
| *prfA*-H4 | UGUAAAAAACAUCAUUUAGCGUGACUUUCUUUCAACAGCUAACAAUUGUUGUUACUCGG GAAUGUUUUUAGGGUAUUUUAAAAAAGGGCGAUAAAAAACGAUUGGGGGAUGAGACAUG AACGCUCAA |
| *prfA*-L5 | UGUAAAAAACAUCAUUUAGCGUGACUUUCUUUCAACAGCUAACAAUUGUUGUUACUGCCU--- UUUUUUAGGGUAUUUUAAAAAAGGGCGAUAAAAAACGAUUGGGGGAUGAGACAUGAACGC UCAA |

a. Mutations are indicated by **red/bold**. Sequence mutations show mutated nucleotide, while deletions are denoted by **-**.
